# Shared gaze reflects shared aesthetic experiences

**DOI:** 10.64898/2026.01.30.702749

**Authors:** Mustafa Alperen Ekinci, Daniel Kaiser

**Author notes:** Correspondence to: Mustafa Alperen Ekinci, **Email:**. **Author Contributions:** M.A.E and D.K. designed research; M.A.E performed research; M.A.E analyzed data; D.K. provided equipment and advice on study design and analysis; M.A.E and D.K. wrote the paper. **Competing Interest Statement:** The authors declare no competing interest.

## Abstract

When individuals view the same visual input, they often differ in their aesthetic appeal judgments, yet why people differ remains largely unclear. Here, we tested whether individual differences in aesthetic experience are linked to differences in visual exploration. In two experiments, participants watched the documentary “Home” while their eye movements were recorded. In Experiment 1, participants continuously rated aesthetic experience throughout the movie, whereas in Experiment 2, they watched the first half without a task and rated aesthetic experience only during the second half. Inter-individual similarity in gaze patterns, assessed using fixation heatmaps across time, predicted similarity in aesthetic appeal judgments in both experiments. Notably, in Experiment 2, gaze similarity during free viewing in the first half of the movie predicted similarity in aesthetic ratings during the second half, indicating that incidental eye movement patterns predict aesthetic experiences. Together, these results show that shared gaze patterns are linked to shared aesthetic experiences under naturalistic, dynamic viewing conditions.

## Introduction

People diverge widely but reliably in their judgments of aesthetic appeal, even when viewing the same stimulus (1, 2), with variation across visual categories and settings (3, 4). Yet the sources of these differences remain poorly understood. Here, we ask whether individual differences in aesthetic appeal are related to differences in visual exploration of naturalistic, dynamic scenes. Eye movements across scenes, faces, and artworks indeed differ across individuals in stable ways (5-7). Although gaze patterns have been linked to aesthetic judgments in various contexts (8), most prior work has focused on group-level effects, leaving the contribution of individual gaze patterns unresolved (7, 9). In this study, we examined whether individual differences in gaze are related to individual differences in aesthetic appeal. In two eye-tracking experiments, participants watched the documentary “Home” while continuously rating their aesthetic appeal (10). By relating participant-to-participant similarities in gaze behavior to similarities in aesthetic appeal ratings, we demonstrate that shared visual exploration is reliably associated with shared aesthetic experiences.

## Results

### Experiment 1

In Experiment 1, thirty participants watched the documentary “Home”, which features a diverse range of pleasing and unpleasing natural scenes from around the world. They watched the documentary in 9 segments in different random orders (mean duration = 505.1 s) while continuously rating its aesthetic appeal using a slider presented throughout the movie (Fig. 1A). From the continuous rating data, we created rating similarity matrices by correlating aesthetic appeal ratings within each segment for each pair of participants and then averaging the matrices across segments.

**Figure 1.**
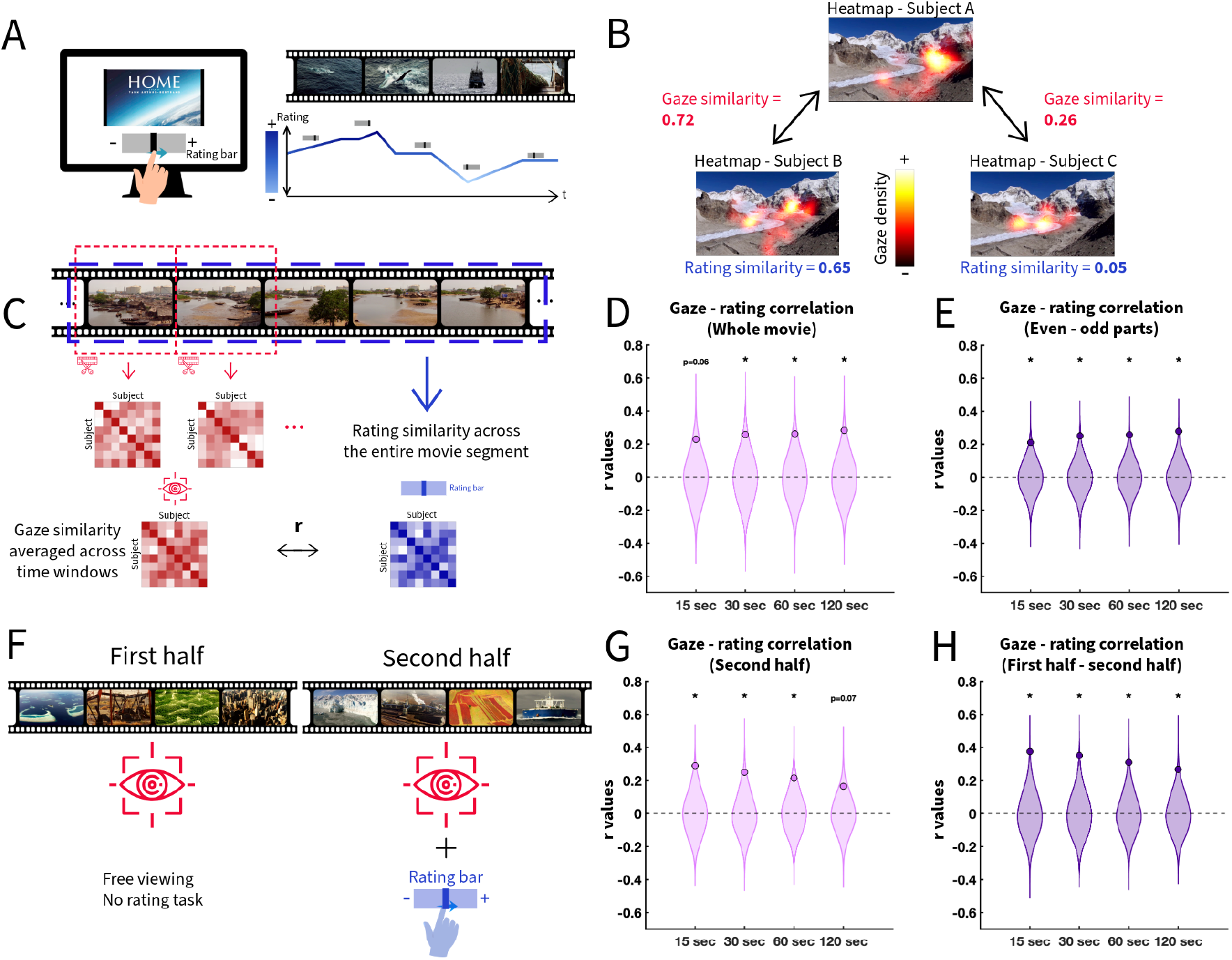
(A) Paradigm. While watching the documentary movie “Home”, participants continuously rated their aesthetic experiences moment-to-moment. (B) Example gaze heatmap similarities and rating similarities for a single movie segment in three participants. (C) Analysis approach. Eye gaze similarity was computed for continuous temporal chunks (15 s, 30 s, 60 s, or 120 s duration) within each movie segment, and a cumulative gaze similarity matrix was obtained by averaging across all available chunks. This cumulative gaze similarity was then correlated with the aesthetic appeal rating similarity to examine whether individual differences in gaze similarity were associated with differences in aesthetic ratings. (D) We observed a robust correlation between inter-subject similarities in gaze patterns and inter-subject similarities in aesthetic appeal ratings. This was true for all chunk sizes (with a marginal effect for the 15s chunks) and (E) also when gaze similarity in odd parts of the movie was correlated to rating similarity in the even parts and vice versa. Dots indicate observed correlations, and shaded areas represent distributions from a permutation test. Asterisks indicate *p*<0.05. (F) In Experiment 2, participants watched the first half of the movie without any task while their eyes were being tracked, whereas in the second half, we surprised them with the aesthetic appeal rating. (G) Focusing on the second half of the movie, the findings from Experiment 1 were replicated, with significant effects for all but the 120s time window. (H) Interestingly, the gaze similarity in the first half (without the rating task) also significantly correlated with the aesthetic ratings in the second half (across all chunk sizes), showing that incidental gaze patterns during free viewing are diagnostic of aesthetic experiences during an explicit task performed on different visual inputs.

We extracted fixations within the movie window as our primary measure of gaze behavior. Within each movie segment, gaze similarity matrices were created for temporal chunk sizes of 15 s, 30 s, 60 s, and 120 s. For each chunk size, we generated fixation heatmaps (Gaussian blur with *σ* = 1°) and computed gaze similarity as the pairwise correlation of these heatmaps across participants, averaged over all chunks and segments (Fig. 1C).

Next, we correlated the lower off-diagonal entries in the rating similarity matrices and gaze similarity matrices (Fig. 1B). Critically, gaze similarity and rating similarity were significantly correlated for the 30 s, 60 s, and 120 s temporal chunk sizes (*r* = 0.25–0.28; *p* < 0.04; permutation tests) with a numerically trending effect for the 15 s chunk size (*r* = 0.22; *p* = 0.06; permutation tests) (Fig. 1D). This relationship also held when gaze similarities were computed for one half of the movie and rating similarities were computed from the other half (*r* = 0.21–0.27; *p* <0.01 for all chunk sizes; permutation tests) (Fig. 1E). Together, this establishes a reliable link between individual differences in gaze and aesthetic appeal.

### Experiment 2

In Experiment 2, participants watched the first half of the movie under free-viewing conditions before we surprised them with the continuous aesthetic appeal rating in the second half (Fig. 1F). This enabled two critical analyses: First, we could replicate the results from Experiment 1, using only the second half of the movie. Second, we could test whether incidental gaze similarities during free viewing in the first half of the movie predict similarities in aesthetic appeal judgments in the second half.

Focusing on the second half of the movie, gaze similarity again predicted rating similarity, across all temporal chunk sizes (*r* = 0.21–0.29; *p* < 0.03; permutation tests) except the 120 s chunk (*r* = 0.17; *p* = 0.07; permutation tests), essentially replicating the previous effects (Fig. 1G). Furthermore, gaze similarity in the first half (without an explicit rating task) also significantly predicted aesthetic ratings in the second half (*r* = 0.31–0.37; *p* < 0.01 for all chunk sizes; permutation tests) (Fig. 1H). Shared patterns of visual exploration are thus linked to shared aesthetic appeal judgments, even when gaze is recorded during free viewing, suggesting that incidental eye movements that occur during free viewing predispose idiosyncrasies in aesthetic experiences.

## Discussion

Our results demonstrate that inter-individual similarity in gaze patterns predicts inter-individual similarity in aesthetic experience, providing a new explanation for idiosyncrasies in aesthetic appeal. Shared gaze patterns, or gaze synchrony, likely reflect shared attentional priorities (11), resulting in similar visual information entering cognitive systems and producing more aligned interpretations and affective responses (7, 9, 12). This account is consistent with evidence linking individual gaze patterns to subjective scene descriptions (13).

Strikingly, similarity in gaze patterns during passive viewing of the first half of the movie predicted similarity in aesthetic appeal ratings for the second half of the movie. While it needs to be determined how different the stimuli for assessing gaze similarities and aesthetic appeal similarities can be for such effects to still emerge, this finding opens up exciting possibilities: Future work could estimate inter-individual similarities in gaze on independent benchmark tasks (14) and use these similarities to predict inter-individual similarities in aesthetic appeal for completely different stimuli, from artworks to design concepts.

Two interesting questions remain open. First, can we attribute a causal role to eye movements in shaping aesthetic experiences? Future work could use cued eye movement sequences or foveal replay of gaze trajectories (15) to test how altered gaze sequences in turn change aesthetic appeal. Second, what is the nature of the divergences in gaze that relate to diverging aesthetic experiences? Candidate explanations are individual tendencies to explore semantic categories (6), individual spatial biases (16), differences in overall saccade dynamics (17), and idiosyncratic ways of maximizing expected sensory reward (18).

Finally, our findings highlight broader implications for understanding shared aesthetic experiences in collective settings such as cinema, museums, or virtual reality. By linking attentional dynamics to aesthetic convergence, our work suggests that gaze similarity could serve as a real-time marker of collective engagement and preference formation.

## Materials and Methods

Thirty participants took part in Experiment 1 (*M*_*age*_ = 26.50, *SD* = 4.25; 16 females), and another thirty in Experiment 2 (*M*_*age*_ = 26.03, *SD* = 4.31; 23 females). The study was approved by the General Ethics Committee of Justus Liebig University Giessen (approval no. AZ25/22), and all participants provided written informed consent before participation. Data and analysis scripts are available on OSF: https://osf.io/qcbx3. Full method details can be found in SI Methods.

## Acknowledgments

This work was supported by the Deutsche Forschungsgemeinschaft (DFG),KA4683/6-1 (project no. 536053998); and under Germany’s Excellence Strategy (EXC 3066/1, “The Adaptive Mind”, project no. 533717223). It was further supported by a European Research Council (ERC) Starting Grant (PEP, ERC-2022-STG 101076057). Views and opinions expressed are those of the authors only and do not necessarily reflect those of the funders. Neither the funders nor the granting authority can be held responsible for them. We thank Tanja John for helping with the data collection.

## Supporting Information Text

### Materials and Methods

#### Participants

The study comprised two different experiments. In Experiment 1, a total of 37 participants with normal or corrected-to-normal vision participated. Data from seven participants were excluded (five due to incomplete data and two due to a lack of variance in the rating task), resulting in a final sample of 30 participants (*M*_*age*_ = 26.50, SD = 4.25; 16 females). In experiment 2, 30 participants with normal or corrected-to-normal vision took part (*M*_*age*_ = 26.03, *SD* = 4.31; 23 females). All participants provided written informed consent and were reimbursed at a rate of €10 per hour for their participation. The study was approved by the General Ethics Committee of Justus Liebig University Giessen (approval no. AZ25/22).

#### Movie Stimulus

In both experiments, participants watched the documentary “Home” divided into nine segments (474.3, 534.8, 529.3, 535.2, 520.7, 537.6, 533.3, 523.3, and 357.7 seconds; mean duration = 505.1 seconds). The movie was presented without sound. The segments did not overlap, and each was cut at the end of a scene to preserve coherence within segments. In Experiment 1, participants watched the movie in one of 8 randomized sequences of segments. The movie has a relatively weak narrative continuity, which is essentially absent without the sound. In Experiment 2, all participants watched the movie in the original segment order.

After collecting data from eight participants in Experiment 1, we slightly improved movie clipping, removing some black screens, which lasted only around 2 minutes in total (3.38% of the total duration). These slight differences between movie versions did not have systematic effects on gaze variability across temporal chunks (correlation of gaze similarity with movie clipping similarity: r = -0.12) and rating variability (correlation of rating similarity with movie clipping similarity; *r* = -0.07). In Experiment 2, the same improved version of movie clipping was used.

#### Procedure

In both experiments, participants were seated in a dark room with their head stabilized using a chin- and forehead rest, positioned 68 cm from the display. Gaze data were acquired using a Tobii Pro Fusion (Nasdaq Stockholm: TOBII, Stockholm, Sweden). Stimuli were presented on a monitor with a resolution of 1280 × 720 pixels and a refresh rate of 100 Hz. At this viewing distance, the stimuli covered 28.7 × 16.4 degrees of visual angle. While watching the movie segments, participants rated aesthetic appeal on a rating scale displayed below the movie, ranging from -144 to +144 in increments of 6 units (49 discrete levels). The experiment was programmed using Psychtoolbox version 3.0.19 (1) in MATLAB R2022a (MathWorks, Natick, MA, USA) on a Windows 10 PC.

The experiment began with a calibration of the eye tracker, which was repeated after every two blocks. Participants were instructed to freely view the movie, keep their heads still, and pay attention throughout the session.

### Data analysis

#### Gaze analysis

Fixations were extracted as the primary measure of gaze behavior. A fixation was defined as a cluster of gaze points lasting at least 200 ms within a 1° radius (2). Only fixations that occurred within the movie frame (28.7 × 16.4 visual degrees) were included in the analysis.For each movie segment, gaze similarity matrices were computed based on temporal chunks of 15, 30, 60, or 120 seconds of continuous viewing, as well as for the entire segment. For each chunk, fixation heatmaps were generated (Gaussian blur with σ = 1°), and gaze similarity was quantified as the pairwise correlation between these heatmaps across participants. The resulting correlations were then averaged across all chunks and segments.

#### Rating analysis

We calculated each participant’s average reaction-time delay in the continuous rating task and temporally shifted their rating data accordingly. To this end, we first identified cuts in the movie by computing frame-to-frame pixel differences, and the average time until participants’ first rating change following each cut was calculated and used to shift the rating data. The shifting was always limited to a maximum of two seconds. Additionally, we excluded the first two seconds of each movie segment since it always took participants more time to adjust their ratings when the movie segment started. We assessed inter-individual rating similarity by computing pairwise correlations of the adjusted ratings across participants. In Experiment 1, to align rating vectors across the original movie clipping (participants 1-8) and slightly trimmed versions (participants 9-30), we resampled the continuous aesthetic rating time series for the pairwise correlations between participants’ ratings.

#### Relating idiosyncrasies in gaze and aesthetic appeal

To examine whether individual differences in aesthetic ratings are linked to individual differences in gaze behavior, we correlated the off-diagonal entries of the rating similarity matrices with those of the gaze similarity matrices across all temporal chunk sizes and the whole movie segment. In Experiment 1, this analysis was conducted over the entire movie. In Experiment 2, the same analysis was applied only to the second half of the movie, during which participants provided aesthetic ratings.

For Experiment 1, we further split the data into even and odd segments. To obtain a cross-validated estimate of the relationship, we repeated the analysis by correlating gaze similarity from the even-numbered movie segments with rating similarity from the odd-numbered segments, and vice versa. Then, we averaged the two resulting correlations.

We further assessed the reliability of individual differences in gaze and aesthetic appeal ratings by correlating the similarity matrices computed from the even-numbered segments with the similarity matrices computed from the odd-numbered segments. The reliability of gaze similarities across temporal chunk sizes was *r* = 0.78 - 0.84, and the reliability of rating similarities was *r* = 0.69.

Finally, in Experiment 2, we examined whether gaze similarity during the first half of the movie, when participants watched the movie under free-viewing conditions, predicted rating similarity in the second half, when aesthetic appeal ratings were collected.

#### Statistical testing

Statistical significance was assessed using a one-tailed permutation test with 10,000 iterations. In each iteration, the order of rows and columns of the rating similarity matrix was randomly permuted to disrupt the correspondence between the rating-based and eye-based similarity structures, thereby generating a null distribution of correlation values expected by chance. The observed correlation was then compared against this null distribution to determine whether it was significantly greater than chance.

**Figure S1.**
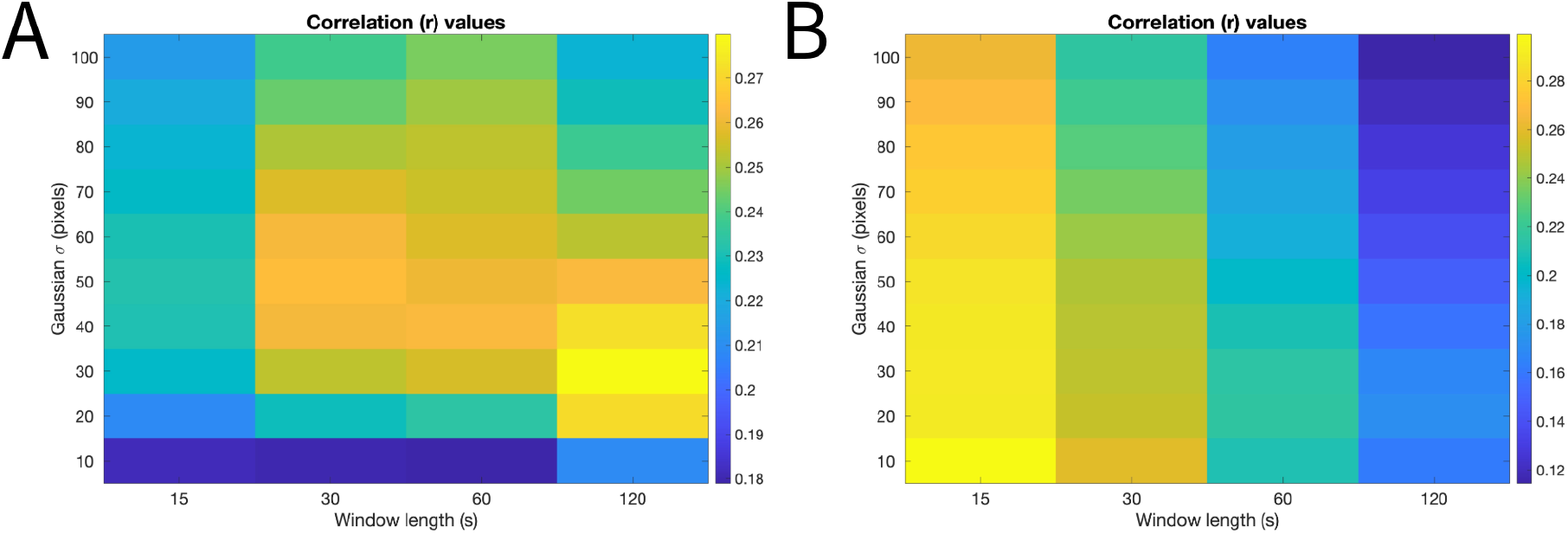
Correlation analyses between gaze similarity and rating similarity were performed across all temporal chunk sizes and heatmap smoothing sigma values in Experiment 1 (A) and Experiment 2 (B). Note that 1 visual degree (32 pixels) was used for the main analysis.

## References

1. E. Martinez, F. Funk, A. Todorov, Quantifying idiosyncratic and shared contributions to judgment. Behav. Res. Methods 52, 1428–1444 (2020).

2. M. Pombo, D. G. Pelli, Aesthetics: It’s beautiful to me. Curr. Biol. 32, R378–R379 (2022).

3. E. A. Vessel, J. Stahl, N. Maurer, A. Denker, G. G. Starr, Personalized visual aesthetics. In Human Vision and Electronic Imaging XIX (SPIE), Vol. 9014, 196–203 (2014).

4. L. Knoll, J. Mikuni, E. Specker, Looking at people looking at art: Observations of art interactions in an everyday urban environment. Front. Psychol. 16, 1658946 (2025).

5. M. S. Castelhano, J. M. Henderson, Stable individual differences across images in human saccadic eye movements. Can. J. Exp. Psychol. 62, 1–14 (2008).

6. B. de Haas, A. L. Iakovidis, D. S. Schwarzkopf, K. R. Gegenfurtner, Individual differences in visual salience vary along semantic dimensions. Proc. Natl. Acad. Sci. U.S.A. 116, 11687–11692 (2019).

7. N. Sharvashidze, A. C. Schütz, Task-dependent eye-movement patterns in viewing art. J. Eye Mov. Res. 13, 12 (2020).

8. W. Chen, R. Ruan, W. Deng, J. Gao, The effect of visual attention process and thinking styles on environmental aesthetic preference: An eye-tracking study. Front. Psychol. 13, 1027742 (2023).

9. P. Francuz, I. Zaniewski, P. Augustynowicz, N. Kopiś, T. Jankowski, Eye movement correlates of expertise in visual arts. Front. Hum. Neurosci. 12, 87 (2018).

10. J. Ortega, D. Whitney, Continuous affect tracking reveals that overestimation during the recollection of affect is idiosyncratic and stable. J. Vis. 25, 14 (2025).

11. B. Bühler et al., On task and in sync: Examining the relationship between gaze synchrony and self-reported attention during video lecture learning. Proc. ACM Hum.-Comput. Interact. 8, 1–18 (2024).

12. E. A. Vessel, N. Maurer, A. H. Denker, G. G. Starr, Stronger shared taste for natural aesthetic domains than for artifacts of human culture. Cognition 179, 121–131 (2018).

13. D. Kollenda, A. S. Reher, B. de Haas, Individual gaze predicts individual scene descriptions. Sci. Rep. 15, 9443 (2025).

14. M. Linka, B. de Haas, OSIEShort: A small stimulus set can reliably estimate individual differences in semantic salience. J. Vis. 20, 13 (2020).

15. J. C. Bush, P. C. Pantelis, X. Morin Duchesne, S. A. Kagemann, D. P. Kennedy, Viewing complex, dynamic scenes “through the eyes” of another person: The gaze-replay paradigm. PLOS One 10, e0134347 (2015).

16. Z. Wang, Y. Murai, D. Whitney, Idiosyncratic perception: A link between acuity, perceived position and apparent size. Proc. R. Soc. B 287, 20200825 (2020).

17. J. M. Henderson, S. G. Luke, Stable individual differences in saccadic eye movements during reading, pseudoreading, scene viewing, and scene search. J. Exp. Psychol. Hum. Percept. Perform. 40, 1390–1400 (2014).

18. Brielmann, P. Dayan, A computational model of aesthetic value. Psychol. Rev. 129, 1319–1337 (2022).

## References

1. Kleiner M, Brainard DH, Pelli DG (2007) What’s new in Psychtoolbox-3? Perception 36(ECVP Abstract Supplement):14.

2. Krassanakis V, Filippakopoulou V, Nakos B (2014) EyeMMV toolbox: An eye movement post-analysis tool based on a two-step spatial dispersion threshold for fixation identification. J Eye Mov Res 7(1):1–10.

